# ATAV: a comprehensive platform for population-scale genomic analyses

**DOI:** 10.1101/2020.06.08.136507

**Authors:** Zhong Ren, Gundula Povysil, David B. Goldstein

## Abstract

**Background:** A common approach for sequencing studies is to do joint-calling and store variants of all samples in a single file. If new samples keep being added or controls are re-used for several studies, the cost and time required to perform joint-calling for each analysis can become prohibitive.

**Results:** We present ATAV, an analysis platform for large-scale whole-exome and whole-genome sequencing projects. ATAV stores variant and per site coverage data for all samples in a centralized database, which is efficiently queried by ATAV to support diagnostic analyses for trios and singletons, as well as rare-variant collapsing analyses for finding disease associations in complex diseases. Runtime logs ensure full reproducibility and the modularized ATAV framework makes it extensible to continuous development. Besides helping with the identification of disease-causing variants for a range of diseases, ATAV has also enabled the discovery of disease-genes by rare-variant collapsing on datasets containing more than 20,000 samples. Analyses to date have been performed on data of more than 110,000 individuals demonstrating the scalability of the framework.

The ATAV data browser (http://atavdb.org/) is a web-based interface that allows users to easily access variant-level data directly from the database. Summary-level data for more than 40,000 samples can be queried by the general public representing a mix of cases and controls of diverse ancestries. Users have access to phenotype categories of variant carriers, as well as predicted ancestry, gender, and quality metrics. In contrast to many other platforms, the data browser is able to show data of newly-added samples in real-time and is therefore evolving rapidly as more and more samples are sequenced.

**Conclusions:** Since all code is freely available on GitHub, ATAV can easily be used by other groups to build up their own platform, database, and user interface. In addition to that users can query one of the largest variant databases for patients sequenced at a tertiary care center and look up their own genes or variants of interest.

## Background

Diagnostic and cohort sequencing studies benefit from the analysis of a large number of samples combined with similarly processed controls. A common approach to reach the necessary scale for analysis is to use a joint-calling procedure and store all samples in a single VCF file[1, 2]. While effective in allowing a single analysis of all samples included in the single VCF file, this approach has significant limitations. Perhaps most importantly, this approach is not amenable to ongoing analyses as new samples become available. Moreover, when projects combine multiple cohorts that were not sequenced together and in which controls might be re-used for several studies, the cost and time required to perform joint-calling for each analysis can become prohibitive. In addition to these considerations, typical sequencing file formats (VCF, BAM) place a sizeable overhead in moving these data from physical storage to the compute nodes for dynamic and multi user analysis needs. Furthermore, standard diagnostic and case-control studies leverage a range of filtering parameters, including variant calling (genotype quality, read coverage), variant annotation (gene, effect), internal population frequencies (minor allele frequency, genotype frequency) and external dataset filters (gnomAD[3], RVIS[4]) to identify “qualifying variants” that meet a specific set of user-defined criteria. These sophisticated needs place an additional burden on creating an audit trail for re-analyses and reproducibility. As the size and number of simultaneous users increase, ad-hoc analyses become prohibitively inefficient in the conventional single joint-genotyped VCF framework.

To address these constraints and dynamic analyses needs, we have developed ATAV (see Figure 1) to streamline genomic analysis needs ranging from the standard diagnostic case interpretation to large-scale cohort analyses for disease-associated gene discovery. The ATAV platform is built on an open source relational database. The database (ATAVDB) is configured with a feature allowing data replication across a cluster of nodes. ATAVDB contains sample variant data, read coverage data, variant annotation data, external annotation data, and metadata. A data pipeline toolkit extracts variants, annotations and associated quality data from VCF files and the coverage and genotype quality from bam files. Currently the Institute for Genomic Medicine (IGM) at Columbia University has data of over 100K whole exomes, and the coding-regions of over 10K whole genomes stored in ATAVDB. It contains over 24 billion variant calls from over 220 million distinct genomic co-ordinates and read coverage information for all samples.

**Figure 1:**
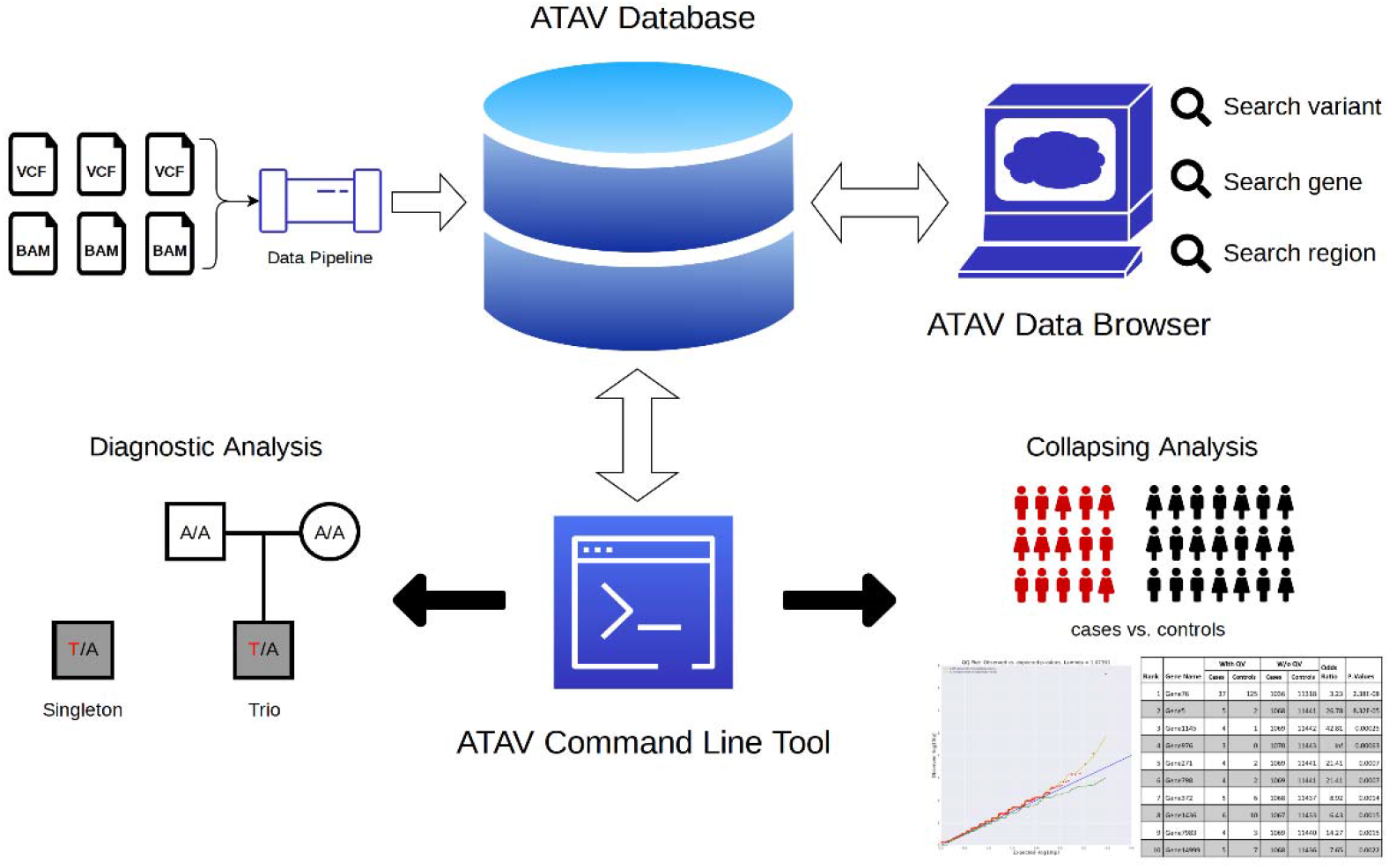
ATAV platform framework overview: Data extracted from single-sample VCF and BAM files is stored in the ATAV database, the ATAV data browser and the ATAV command line tool retrieve information from the ATAV database for variant look-up, diagnostic analyses and association studies using rare-variant collapsing.

## Implementation

### Database

We use Percona Server for MySQL and its high-performance storage engine Percona TokuDB to improve scalability and operational efficiency. In the database, we store a universal variant list across all samples, annotation data that is annotated through ClinEff[5], sample variants calls and associated quality metrics, as well as all site’s coverage data for inferring reference alleles at non-call sites. In addition to that, ATAVDB stores external databases such as allele frequencies from gnomAD[3], ExAC[6], or, DiscovEHR[7], scores such as GERP++[8], TraP[9], LIMBR[10], MTR[11], RVIS[4], subRVIS[12], REVEL[13], PrimateAI[14], CCR[15], as well as clinical annotations from ClinVar[16, 17], ClinGen[18], HGMD[19], and OMIM (see Figure 2). ATAV has an external data plugin code structure, which allows quick code integration of gene based, site based and variant based data.

**Figure 2:**
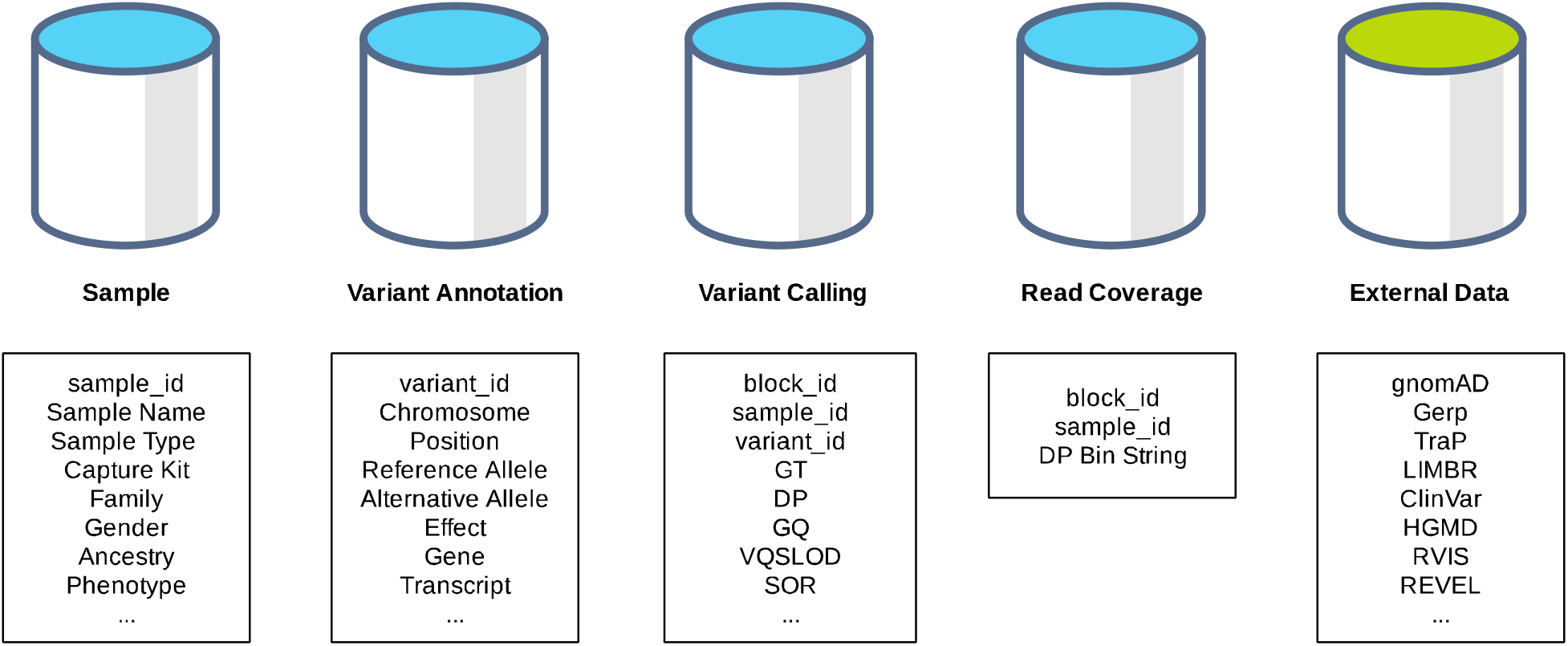
ATAV core database schema and external databases

For efficiently storing coverage information for every site and every sample, the ATAV data pipeline parses through the bam files to generate read coverage data and converts site coverage values into bin values: a [0-9]; b [10-19]; c [20-29]; d [30-49]; e [50-199]; f >=200. A run-length encoding procedure is used to further compress data within fixed 1000 bp block regions (see Figure 3). This way the data size is reduced by about 1000 times making it possible to store the coverage information for more than 100K samples. The information that has been loaded into ATAVDB has been determined over many years of applied use to be that information that is most often required for the standard genetic analyses performed as part of both diagnostic genetic studies and gene discovery. For example, in diagnostic analyses for identifying *de novo* mutations in affected children, it is necessary to know that the parental samples have sufficient coverage at the relevant site, but not necessary to know the precise number of reads, leading to the binning strategy on coverage described above. For the vast majority of applications, we have found that the necessary information can be economically stored and retrieved as described.

**Figure 3:**
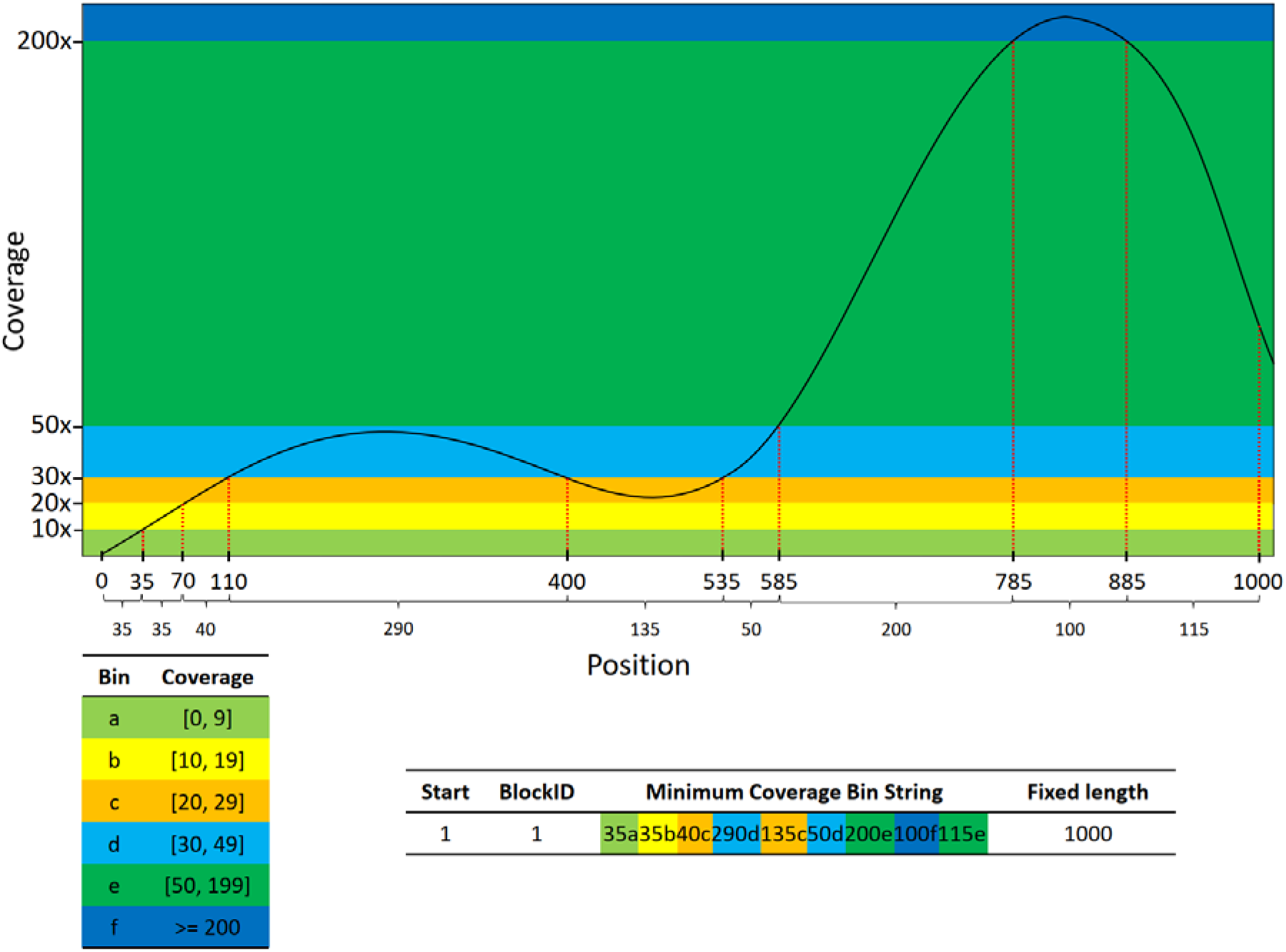
Efficient storage of coverage information: The per site coverage value is converted into a fixed 1000 base pair length bin string by first converting coverage values into bins (a-f) and then using a run-length encoding procedure to further compress data within fixed 1000 bp block regions by summarizing consecutive coverage values within the same bin.

### Platform architecture

The platform architecture is depicted in Figure 3. Users login to the head node to run ATAV jobs which will automatically allocate resources and submit jobs to the cluster. A standard setup with a 6 node Sun Grid Engine (SGE) cluster (2×10 Cores, 128GB RAM) allows the running of at least 100 jobs at once. Each job will query a slave database with minimum database connections. Using a local customized bioinformatics pipeline, it is possible to continue loading new samples into the master database which will automatically replicate to all slave databases.

**Figure 4:**
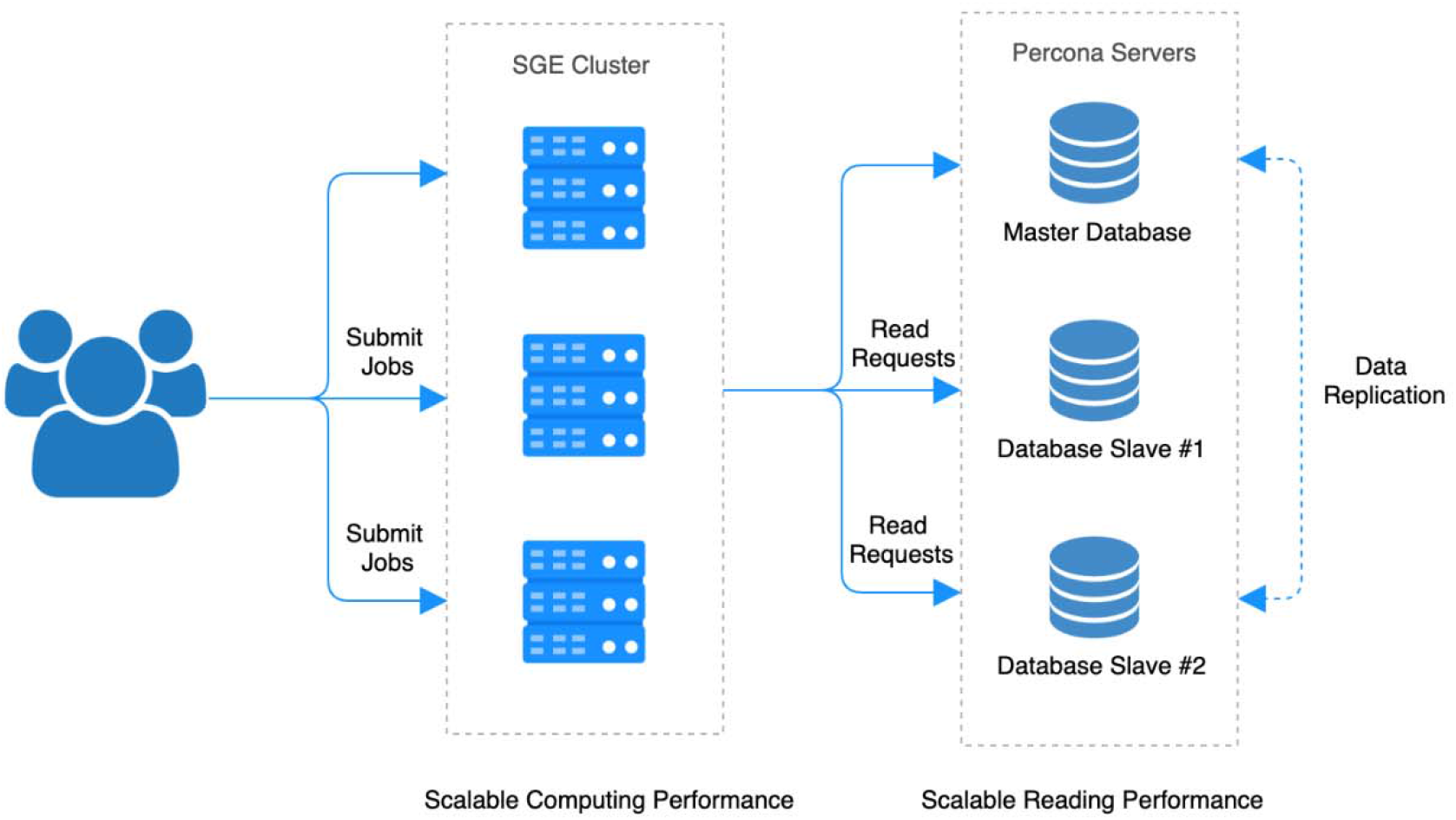
Platform architecture: The user submits jobs to an SGE cluster, the ATAV job will then query to get data from a replicated slave database.

### Application

The ATAV command line tool is the interface to ATAVDB. Written in java, ATAV consists of three modules. (i) The command line parser and query engine translate user defined parameters and the input sample list (in PLINK’s ped format[20]) into an efficient SQL query for interrogating the relational database, (ii) A runtime variant object creator parses SQL output into a collection of variant objects. Each variant object includes variant information (genomic coordinates, annotation), variant calls in sample list, sample genotype calls at co-ordinates without a called variant and external annotation data. (iii) A statistical analyses module iterates over the variant objection collection to perform downstream analyses. ATAV currently supports tests for diagnostic analyses such as identifying putative *de novo* and inherited genotypes of interest in trios, and a framework for performing region-based rare-variant collapsing analyses that identify genes or other genomic units that carry an excess of qualifying variants among cases in comparison to the background variation observed in internal controls of convenience in ATAVDB.

The modularized ATAV framework makes it extensible to continuously develop new functions that operate on sequencing/variant data sets. Critical to data integrity, all ATAV analyses allow an auditable log of software and database version, filter parameters adopted, the input sample lists used in the specific run and the runtime logs that ensure full reproducibility.

The analysts and researchers of the IGM, have run about 26,000 ATAV job within the last year. 15,000 jobs completed in minutes, 7,000 jobs completed in hours, and the remaining 4,000 jobs completed within two days.

The ATAV data browser is a web user interface that allows everyone to access variant-level data directly from the full data set (authorized user) or public available data set (anonymous user) in the ATAVDB. It supports the search of variants by gene, region and variant ID. The gene or region view displays a list of variants with allele count, allele frequency, number of samples, effect, gene etc. The variant view (see Figure 5) displays a set of annotations (effect, gene, transcript, PolyPhen[21]) and details about variant carriers (gender, predicted ancestry, phenotype, and quality metrics). It includes links to other public data resources such as Ensembl, gnomAD[3], ClinVar[16, 17] etc. and directly integrates additional annotations via APIs (e.g. the Genoox Franklin API for clinical variant interpretation). The data browser has advanced filters such as a maximum allele frequency threshold to only search rare or ultra-rare variants, restriction to high quality variants or restriction to a certain phenotype. The public view currently contains more than 40,000 samples representing a mix of cases and healthy controls of diverse ancestries. Users can look up potential disease causing variants and see if the phenotype of variant carriers in ATAVDB matches their phenotype of interest. In contrast to many other platforms, the data browser is able to show data of newly added samples in real-time and is therefore evolving rapidly as more and more samples are sequenced.

**Figure 5:**
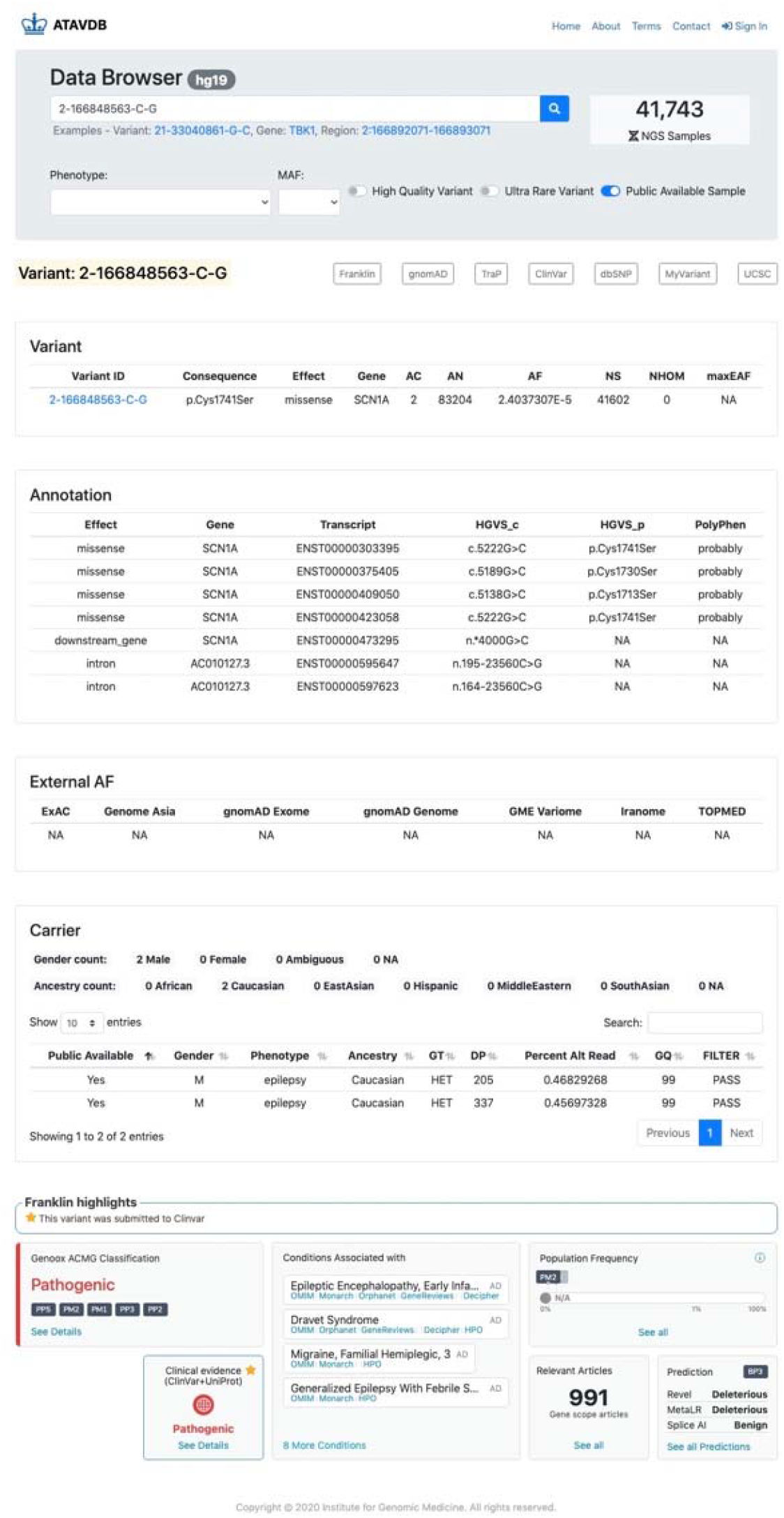
Example of the variant view of the ATAV data browser.

### Analysis

ATAV provides functions for all recommended steps of the rare-variant collapsing workflow recently summarized in Povysil et al. 2019[22]. For the sample pruning steps ATAV creates the necessary input files by pulling data out of ATAVDB and automatically calls existing standard tools such as KING[23], Eigenstrat[24], or FlashPCA[25]. Since the coverage information for every sample and site is already efficiently stored in ATAVDB, ATAV can efficiently compare coverage between cases and controls and provides two different tests to perform coverage harmonization: sites can be removed if cases and controls show differing proportions of individuals with enough coverage[26]; or if a binomial test shows that the case/control status and coverage are not independent[27]. The outputs of the sample pruning and coverage harmonization steps can be used as inputs for dominant or recessive collapsing models. Within the collapsing model call, ATAV selects qualifying variants (QVs) that pass filters based on variant quality (Phred quality (QUAL), genotype Phred quality (GQ), quality by depth (QD), mapping quality (MQ) and variant quality score log-odds (VQSLOD)), variant annotation (effect, pathogenicity prediction scores, intolerance scores), as well as internal and external minor allele frequencies (MAFs). All QVs are used for building the collapsing matrix, a gene-by-individual indicator matrix with a 0 if there is no qualifying variant found in that gene in that individual, and a 1 if there is at least one. This collapsing matrix is used for looking for associations between genes with QVs and the phenotype of interest by using a Fisher’s exact test or Firth-based logistic regression. Finally, quantile-quantile (QQ) plots are created and the genomic inflation factor lambda is estimated using a permutation-based expected distribution of p-values.[26]

A standard collapsing analysis usually consists of several different models that all capture specific types of QVs. While quality control (QC) filters are used for all models, other filters, such as the predicted variant effects or population allele frequencies, depend on the specific model in use. In order to speed up computation, ATAV provides the option of running a general collapsing model first using the QC filters all models have in common and relaxed allele frequency thresholds. The output of this initial model can be used as input for a collapsing-lite function that makes it possible to run the individual collapsing models within minutes since additional filters can just be applied to the previous output and the variant database does not have to be queried again.

All annotations and filters mentioned previously such as QC filters or internal or external MAFs are also important for diagnostic analyses especially for singletons where we cannot use additional family information. In addition to that, ATAV provides special functions for trios and families to reduce the number of potential disease-causing variants in the final output. ATAV leverages information about family structure and affectedness status that is provided by the sample file (PLINK-style ped file). Multiple families can be analyzed at once and related controls are for example automatically removed when calculating control frequencies. Furthermore, the affectedness status is used to decide whether to look for inherited or *de novo* variants.

In the standard trio case of one affected offspring and unaffected parents, ATAV uses a series of functions to extract *de novo* variants, newly compound-heterozygous or newly homozygous variants. For distinguishing compound-heterozygosity from variants that are in-phase, ATAV checks that both parents carry one of the qualifying variants. ATAV not only considers the genotype of the individuals, but also their coverage. If the coverage at a variant site is below a minimum threshold of 10 for any of the individuals the variant is still included in the output, but flagged as possibly *de novo*, possibly newly compound-heterozygous or possibly newly homozygous. Furthermore, ATAV identifies putative parental-mosaic variant transmissions. For each parent-child pair, it extracts all variants that were transmitted from parent to child where the variant in the parent has a low proportion of alternate alleles indicating mosaicism.

ATAV also leverages an external annotation dataset called KnownVar, which combines information from multiple variant and disease databases (e.g. ClinVar[16, 17], HGMD[19], OMIM, ClinGen[18]). The data is stored in ATAVDB and regularly updated. KnownVar annotations are not only included if the “exact” variant has been reported before, but also if a different variant at the same site has been linked to disease. Typical annotations include the associated disease, ClinVar clinical significance, HGMD Class and Pubmed IDs of relevant papers. In addition to that, disease associated variants in close proximity are extracted from HGMD and ClinVar. On a gene level, annotations include the total number of likely pathogenic or pathogenic variants of each category (copy number variation, small insertion/deletion, splice, nonsense, missense) in ClinVar, disease associations and inheritance from OMIM and dosage sensitivity from ClinGen. All the information provided by KnownVar can be used as additional information in the diagnostic setting to evaluate whether a variant can be considered as diagnostic for a specific patient.

## Results

The collapsing framework of ATAV has enabled the confirmation of known and the discovery of novel genes in a wide range of diseases such as epilepsies[28, 29], sudden unexplained death in epilepsy[30], congenital kidney malformations[31], chronic kidney disease[32], amyotrophic lateral sclerosis[33, 34], Alzheimer’s disease[27], retinal dystrophy[35], and idiopathic pulmonary fibrosis[26]. Furthermore the diagnostic framework has helped to identify both diagnostic genotypes in known genes and candidate genotypes in novel genes in a wide range of diseases including stillbirth[36], rare undiagnosed genetic disorders[37, 38], epilepsies[39–41], alternating hemiplegia of childhood[42], and chronic kidney disease[43].

## Conclusions

We present ATAV as an analysis platform for large-scale whole-exome and whole-genome sequencing projects. The most challenging aspect of the initial use of ATAV is that it needs to be used with ATAVDB and requires establishing a similarly structured database and loading into it the necessary data for retrieval. The advantages of the ATAV framework, however, are that 1) it allows continuous real-time analyses of all samples loaded into the database without the need for computationally demanding joint calling preceding each analysis and 2) it allows convenient tracking of precise analyses performed. The newly added ATAV data browser provides easy access even to users with little computational experience by providing an intuitive web interface to query variant-level data directly from the database.

Our experience with this platform on a database carrying more than 100,000 samples indicates that a relational database can be optimized in a way that makes it possible to analyze current large-scale genomic datasets. Our current data processing and storage framework is robust and flexible when combining data from multiple projects and mixing exomes and genomes. ATAV supports diagnostic analyses for trios and singletons, as well as rare-variant collapsing analyses for finding disease associations in complex diseases. Further optimizations are possible such as database sharding which is a horizontal partition of data in a database or search engine. Other potential solutions include storing the data in HDFS (Hadoop Distributed File System) and utilizing Apache Spark to do distributed cluster computing. This would allow the processing of large amounts of variant data in parallel at once speeding up computations and enabling an even further increase in sample sizes.

The goal of ATAV is to work towards standardizing and optimizing storage and data processing for large scale sequencing data across multiple studies and to provide an easy to use interface for users with little computational experience while ensuring full reproducibility.

All code for building ATAV is publicly available, providing a convenient way for other groups to build up their own analysis platform, database, and user interface. In addition to that, since we provide general access to part of our database via the ATAV browser, users can also query one of the largest variant databases available for patients sequenced at a tertiary care center, Currently, the general public can query summary-level data for more than 40,000 samples, but since data of newly sequenced samples are added in real-time this number grows steadily, increasing the value of the database even further as more and more samples are sequenced.

## Availability and requirements

Project name: ATAV

Project home page: https://github.com/nickzren/atav

Operating system(s): Platform independent

Programming languages: Java, Python, R, HTML and Javascript

Other requirements: Java 1.8 or higher, Percona Server 5.6 or higher

License: MIT License

Any restrictions to use by non-academics: No restrictions

## List of abbreviations

VCF: Variant Call Format
BAM: Binary Alignment Map
gnomAD: The Genome Aggregation Database
RVIS: Residual Variation Intolerance Score
ATAVDB: ATAV Database
IGM: Institute for Genomic Medicine
ClinEff: Clinical Variant Annotations Software
ExAC: The Exome Aggregation Consortium
GERP: Genomic Evolutionary Rate Profiling
TraP: The Transcript-inferred Pathogenicity
LIMBR: The Localized Intolerance Model using Bayesian Regression
MTR: The missense tolerance ratio
HGMD: The Human Gene Mutation Database
OMIM: Online Mendelian Inheritance in Man
SQL: Structured Query Language
PolyPhen: Polymorphism Phenotyping
QC: Quality Control
QVs: Qualifying Variants
QUAL: quality score
GQ: genotype quality score
QD: quality by depth score
MQ: mapping quality score
VQSLOD: variant quality score log-odds
MAF: Minor Allele Frequencies
HDFS: Hadoop Distributed File System

## Declarations

### Ethics approval and consent to participate

Not Applicable.

### Consent for publication

Not Applicable.

### Availability of data and materials

The ATAV data browser is hosted at http://atavdb.org/. All code is freely available on GitHub (ATAV command line tool: https://github.com/nickzren/atav, ATAV data browser: https://github.com/nickzren/atavdb).

### Competing interests

D.B.G. is a founder of and holds equity in Praxis, holds equity in Q-State Biosciences, serves as a consultant to AstraZeneca, and has received research support from Janssen, Gilead, Biogen, AstraZeneca, and Union Chimique Belge (UCB). Z.R. and G.P. declare no competing interests.

### Funding

This project was funded by the Institute for Genomic Medicine, Columbia University Irving Medical Center.

### Authors’ contributions

ZR authored the code for ATAV; All authors wrote the paper and revised it. All authors read and approved the final manuscript.

## Acknowledgements

We thank Slavé Petrovski for contributing to the original design of the analysis framework and Quanli Wang for contributing to the original platform development.

## Web links and URLs

ATAV command line tool, https://github.com/nickzren/atav

ATAV data browser, https://www.atavdb.org/

ClinEff, http://www.dnaminer.com/clineff.html

ClinGen, https://clinicalgenome.org/

ClinVar, https://www.ncbi.nlm.nih.gov/clinvar/

Ensembl GRCh37, https://grch37.ensembl.org/

ExAC, http://exac.broadinstitute.org/

dbSNP, https://www.ncbi.nlm.nih.gov/snp/

Franklin, https://franklin.genoox.com/

Iranome, http://www.iranome.com/

MyVariant, http://myvariant.info/

Genome Asia, https://browser.genomeasia100k.org/

GME Variome, http://igm.ucsd.edu/gme/

gnomAD, https://gnomad.broadinstitute.org/

HGMD, http://www.hgmd.cf.ac.uk/ac/index.php

OMIM, https://www.omim.org/

TOPMed hg19, https://bravo.sph.umich.edu/freeze3a/hg19/

TraP, http://trap-score.org/

RVIS, http://genic-intolerance.org/

UCSC Genome Browser, https://genome.ucsc.edu/index.html

## Notes

### Summary of Updates

Added ATAV platform framework overview figure

